# ARP2/3- and resection-coupled genome reorganization facilitates translocations

**DOI:** 10.1101/2021.10.22.465487

**Authors:** Jennifer Zagelbaum, Allana Schooley, Junfei Zhao, Benjamin R. Schrank, Elsa Callen, Shan Zha, Max E. Gottesman, André Nussenzweig, Raul Rabadan, Job Dekker, Jean Gautier

**Author notes:** The University of Texas MD Anderson Cancer Center, Houston, Texas 77030. These authors contributed equally to this work (Allana Schooley, Junfei Zhao).

## Abstract

DNA end-resection and nuclear actin-based movements orchestrate clustering of double-strand breaks (DSBs) into homology-directed repair (HDR) domains. Here, we analyze how actin nucleation by ARP2/3 affects damage-dependent and -independent 3D genome reorganization and facilitates pathologic repair. We observe that DNA damage, followed by ARP2/3-dependent establishment of repair domains enhances local chromatin insulation at a set of damage-proximal boundaries and affects compartment organization genome-wide. Nuclear actin polymerization also promotes interactions between DSBs, which in turn facilitates aberrant intra- and inter-chromosomal rearrangements. Notably, BRCA1 deficiency, which decreases end-resection, DSB mobility, and subsequent HDR, nearly abrogates recurrent translocations between AsiSI DSBs. In contrast, loss of functional BRCA1 yields unique translocations genome-wide, reflecting a critical role in preventing spontaneous genome instability and subsequent rearrangements. Our work establishes that the assembly of DSB repair domains is coordinated with multiscale alterations in genome architecture that enable HDR despite increased risk of translocations with pathologic potential.

## Main

Eukaryotic nuclei are organized into functional domains enriched in proteins and factors involved in nuclear transactions including RNA splicing, transcription, DNA replication, and repair^1^. Repair domains assemble in biomolecular condensates mediated by mechanical forces and multivalent interactions between proteins and nucleic acids^1^. We have shown that the actin nucleator, ARP2/3, and its activator WASP promote clustering of DSBs into homology-directed repair (HDR) domains, which stimulates repair by facilitating DNA end-resection^2^, the initial step of HDR^3^. In turn, resection leads to increased DSB mobility^2^. Thus, HDR domains arise from the coordinated action of actin forces and repair reactions.

Besides facilitating repair^2,4-8^, the impact of ARP2/3 and DSB-induced motion on genome organization is poorly understood. Moreover, the pathologic consequences of assembling DSBs into nuclear domains are not fully known. Imaging of DSBs in mammalian cells has revealed that translocating breaks are mobile^9^, and clustering of DSBs in yeast promotes rearrangements^6,10^. Furthermore, CtIP-dependent resection facilitates translocations in mouse cells^11^, highlighting that DSB end-processing can lead to misrepair. Nevertheless, it remains unclear how proteins that mediate HDR domain formation, including actin nucleators and resection machinery such as the tumor suppressor BRCA1, affect chromosome translocations.

### DNA damage induces local and global chromatin reorganization

DNA damage activates the local, stepwise recruitment of repair proteins to damage sites as well as protein modifications that can spread over megabases along chromatinized DNA, such as the phosphorylation of the histone H2A variant, H2AX^12-14^. Live-cell imaging and clustering analyses of DNA repair foci demonstrate that these processes translate into the 3D reorganization of chromatin^2,15^ but do not fully characterize the genomic features of repair domains and the rules that govern their assembly. To assess the impact of DSBs on genome organization, we performed chromosome conformation capture (Hi-C) in mouse embryonic fibroblasts (MEFs)^16^ and human osteosarcoma cells (U2OS)^17^ harboring an inducible AsiSI restriction endonuclease. These cells express AsiSI fused to a truncated estrogen receptor that translocates to the nucleus upon induction with 4-hydroxy-tamoxifen (4OHT). There are over one thousand AsiSI recognition motifs in the mouse genome. However, cleavage efficiency, as measured by END-seq spike-in assays, revealed that a significant proportion of these sites are not cut in MEFs (**Extended Data Fig. 1a**). Therefore, for quantitative analysis of Hi-C data, we focused on a subset of 97 frequently cut AsiSI sites which showed the highest END-seq signal above background and collectively account for the approximately 100 DSBs per cell^16^. Cells were treated with the ARP2/3 inhibitor, CK-666, for six hours following induction of DSBs (+4OHT). After ensuring that CK-666 did not affect cutting efficiency of AsiSI-ER or the accumulation of damage (**Extended Data Fig. 1b,c**), two biological replicates were performed. Replicates of Hi-C experiments showed comparable phenotypes (**Extended Data Fig. 1d**), and data was pooled for the main figure sets.

Hi-C studies have revealed that the genome can be split into A (open) and B (closed) chromatin compartments that preferentially self-interact and represent open and closed chromatin, respectively^18^. We first sought to examine the impact of DSBs and nuclear actin polymerization on A/B compartment distribution using eigenvalue (principal component, PC1) decomposition of contact matrices^18,19^. Strikingly, eigenvector analysis of intra-chromosomal interactions revealed that following induction of damage, a significant fraction of 250 kb bins, particularly those with eigenvalues closer to zero, had a relative increase in EV1 and apparent flip from B (closed chromatin) to A (open chromatin) (**Fig. 1a; Extended Data Fig. 2a,b**). Approximately 15% (all sites) or 25% (top 97 sites) of B chromatin bins flipped to the A compartment genome-wide or within 2 Mb of frequently digested AsiS1 sites, respectively (**Fig. 1a**). These changes resulted in genome-wide enrichment of open chromatin with a nearly 10 percent increase in the A compartment following damage (**Extended Data Fig. 2a**). An example of a B-to-A compartment flip (blue to red) coinciding with an AsiSI site on chromosome 2 is shown (**Fig. 1b**, DSB #50). Switches were also found at a distance from the AsiSI site, as seen 2 Mb downstream of DSB #8 (**Fig. 1b**). Notably, ARP2/3 inhibition with CK-666, dampened damage-induced compartment flips, indicating a role for nuclear actin polymerization in genome compartmentalization (**Fig. 1a; Extended Data Fig. 2a,b**). Compartment flips during transcriptional regulation correlate with changes in histone modifications^20^. Similarly, restriction endonuclease damage induces megabase-sized chromatin remodeling events, including H2AX phosphorylation, ubiquitin accumulation, and histone H1 depletion^14^, which may contribute to the compartment switching events seen here. We next visualized genome-wide compartmentalization using saddle plots, which display interaction frequencies between pairs of 250 kb loci arranged according to their first eigen values (**Extended Data Fig. 2c**). Compartment strength was calculated using (AA+BB)/(AB+BA) to assess the preference for homotypic (A-A, B-B) over heterotypic (A-B) interactions^21^. Saddle plots and strength quantification revealed that damage increased both homotypic interactions and compartment strength in the B compartment (**Extended Data Fig. 2c**). Notably, ARP2/3 inhibition with CK-666 slightly increased B-B interaction strength in both the presence and absence of damage, further suggesting a role for actin nucleation in compartmentalization (**Extended Data Fig. 2c**).

**Figure 1.**
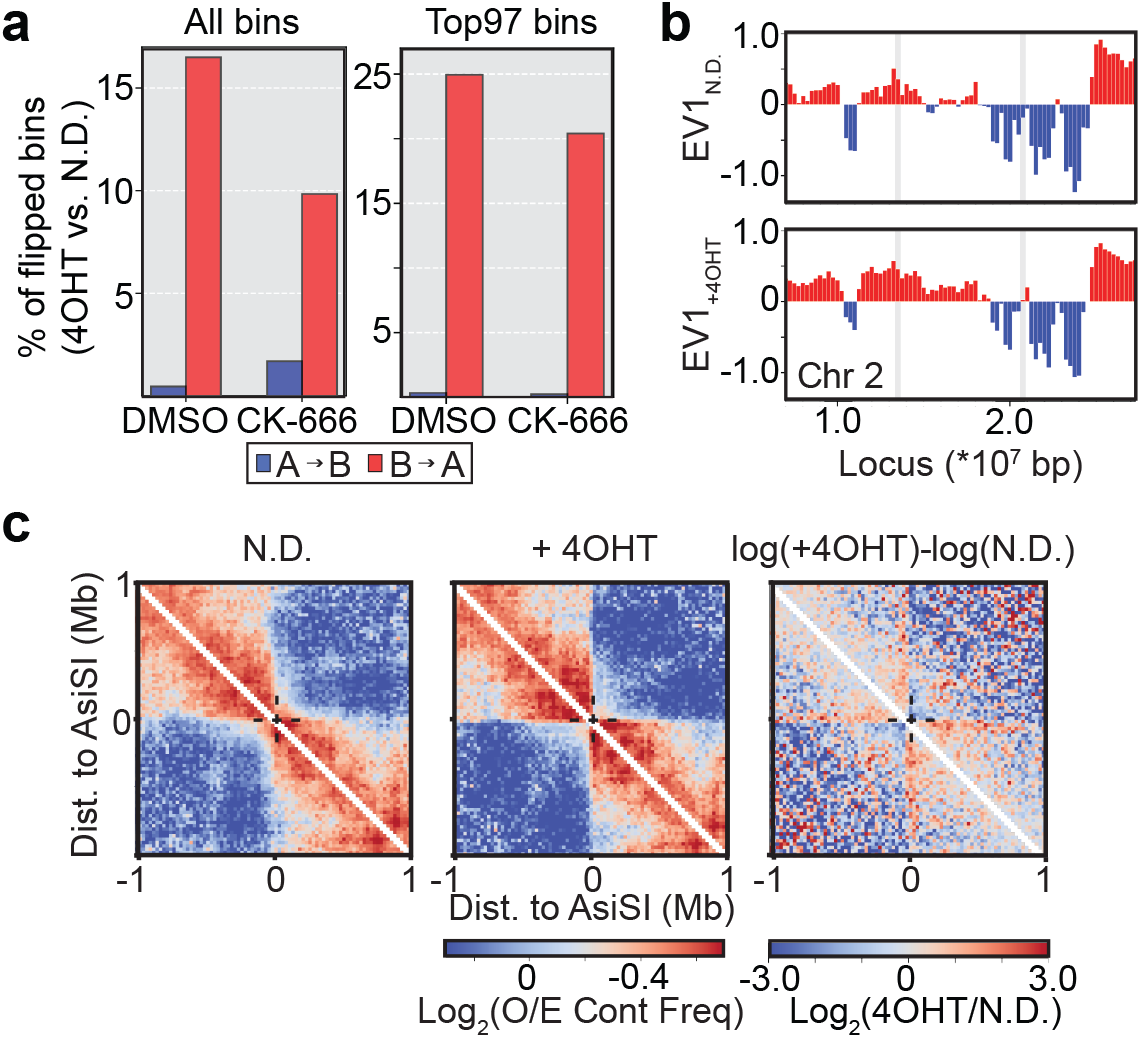
DNA damage induces multiscale alterations of the 3D genome. **a**, Percent of A (open) or B (closed) compartment bins (250 kb) that flip identity genome-wide (left) upon induction of damage with 4OHT genome-wide and for the 2 Mb regions surrounding the top 97 frequently cut AsiSI sites in MEFs - (right). **b**, Representative trajectory of compartment flipping events. First eigenvector (EV1) tracks for *cis* interactions (250 kb bins) normalized to observed-expected. Values are phased by gene density (Active chromatin/A compartment>0, red). Frequently cut AsISI sites are highlighted in grey. **c**, Average log_2_(observed/expected) Hi-C interaction frequency maps of the aggregated 2 Mb regions surrounding the most frequently cut AsiSI sites binned at 25 kb resolution.

Contact probability *P* plotted as a function of genomic distance *s* (*P(s)*) can reveal properties of chromatin architecture including the size and density of cohesin-dependent loops^22^ (**Extended Data Fig. 2d**). The derivative of *P*(*s*) typically displays a local peak at *s* ∼100 kb, corresponding to the average size of loops, followed by a valley at *s* ∼2 Mb. The depth of the valley is related to loop density^22^. Analysis of the derivative of *P*(*s*) for Hi-C data obtained from cells with induced DNA damage revealed a more pronounced valley at *s* ∼2 Mb (**Extended Data Fig. 2d**). This can be interpreted as a general increase in loop density. An increased number of loops/kb of DNA upon damage is consistent with previous observations of increased cohesin recruitment to DSBs, including those induced by restriction endonucleases, that could reflect cohesin-driven loop extrusion at DSBs^23-25^, or possibly more generally genome-wide. Indeed, average Hi-C interaction frequency aggregated at CTCF sites showed increased line formation (**Extended Data Fig. 2e**). Such lines reflect increased cohesin-mediated loop formation where one base of the loop is anchored at the CTCF sites. Hi-C interaction frequency between pairs of convergent CTCF-CTCF sites also increased upon DNA damage (**Extended Data Fig. 2f**). Interestingly, the difference in *P(s)* upon DNA damage, as well as average insulation and loop strength at and between CTCF sites was not diminished in the presence of CK-666 (data not shown).

Next, we aggregated contact matrices spanning 2 Mb around frequently cut AsiSI sites to visualize interactions in the absence (No Damage, N.D.) and presence (+4OHT) of DSBs (**Fig. 1c**). Average contact maps revealed a striking level of organization in undamaged cells, as seen by strong insulation at the AsiSI motifs (**Fig. 1c, left panel**). Importantly, gene set-enrichment analysis (GSEA), which identifies statistically significant relationships between biological states, showed that frequently cut AsiSI sites are enriched in transcriptionally active areas (**Extended Data Fig. 2g**). Upon induction of DSBs, average insulation at these sites increased in both MEFs (**Fig. 1c, middle and right panels**) and in AsiSI-U2OS cells (**Extended Data Fig. 2h**) indicating that this is a conserved feature. This increase is likely driven by a subset of DSBs located at insulated boundaries, or compartment boundaries. This suggests that DNA damage strengthens pre-existing genome organization into more robust DNA repair domains^15,25,26^.

These data reveal multiscale changes in the 3D genomic landscape following damage, including compartment switching that favors open chromatin states, increased loop density, and reinforcement of pre-existing chromatin organization surrounding DSBs. It also suggests that long-range genome reorganization, such as damage-induced compartment flipping that enriches for A compartment, is regulated in part by ARP2/3-driven nuclear actin polymerization.

### Chromosome translocations occur at sites of DSB clustering

In yeast and mammalian cells, DSB mobility drives clustering of DSBs into repair factories^2,4-8^. Therefore, we sought to characterize long-range intrachromosomal interactions using aggregate peak analysis (APA)^27,28^. We piled-up all possible pairwise interactions (304) occurring within chromosomes (in *cis*) between the most frequently cut AsiSI sites. Following damage, we visualized distant DSBs coming together in both MEF and U2OS cells (**Fig. 2a, Extended Data Fig. 3a-d**). DSB cluster enrichment scores were calculated by comparing the average signal intensity at the center of the plot with that of the surrounding area, allowing for relative quantification of interaction frequency between AsiSI sites. Clustering increased upon addition of 4OHT (1.24 to 2.01), indicating that DNA damage triggered increased interactions between distant DSBs within individual chromosomes (**Fig 2a**; **Extended Data Fig. 3c**). These distant DSB-DSB interactions were partially reduced by CK-666 treatment (2.01 to 1.80). These data suggest that this DNA damage-dependent clustering is driven in part by ARP2/3-dependent nuclear actin polymerization.

**Figure 2.**
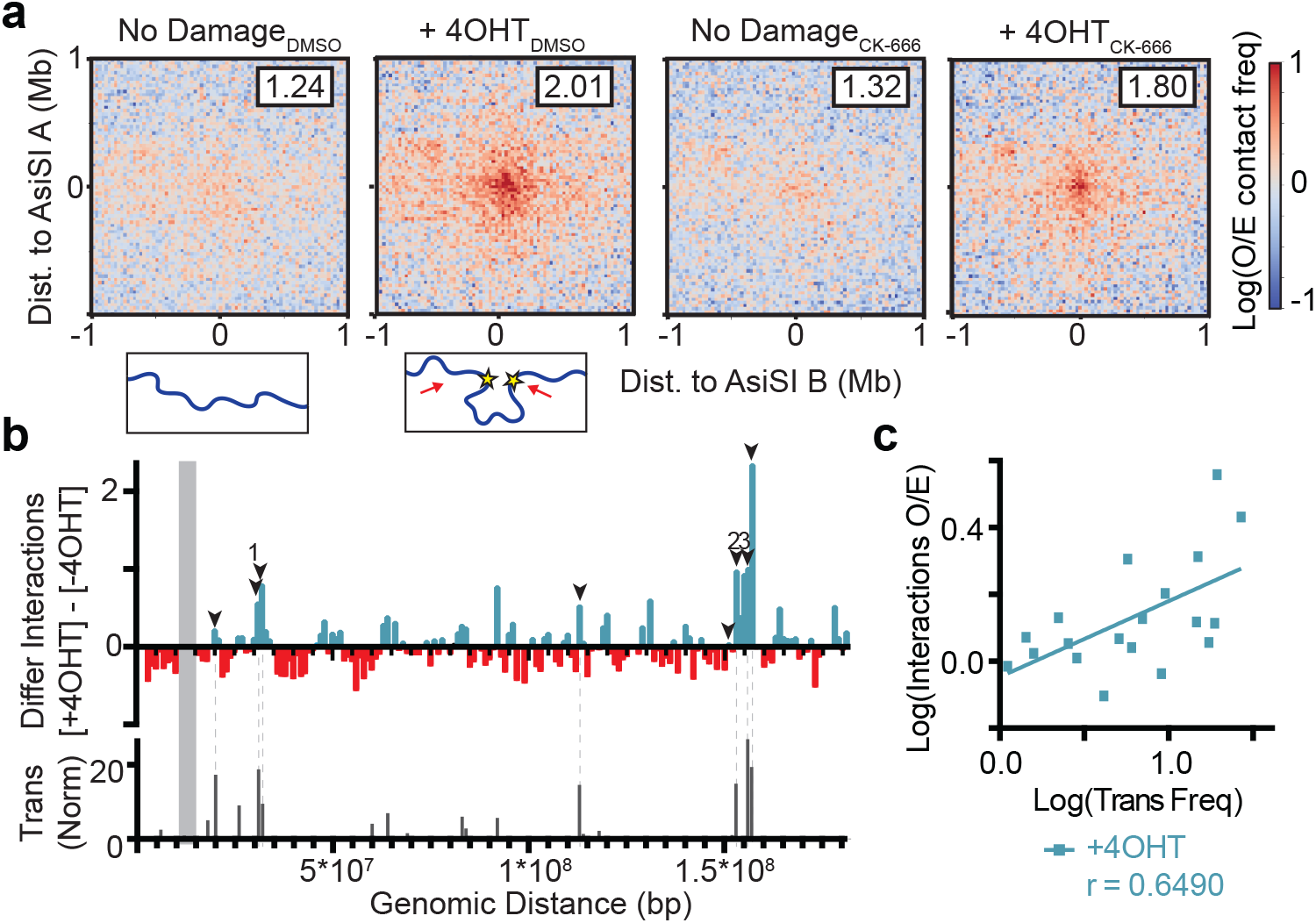
Translocations occur at sites of DSB clustering. **a**, Top: Aggregate peak analysis (APA) displaying the contact frequencies of all possible pairwise combinations of the the top 97 AsiSI digested sites *in cis* (304 interactions between damaged bins). Data is binned at 25 kb and averaged for a 2 Mb flanking window. Log_2_(observed/expected) Hi-C maps are shown in the presence or absence of damage (4OHT) +/-CK-666. For each APA plot, a cluster enrichment score is calculated using the ratios of the average interaction frequency of the 9 central bins (125 kb) / average interaction frequency of the outside bins (125 kb – 1 Mb). Bottom: Schematic visualization of intra-chromosomal interactions and clustering following damage. **b**, Top: Differential interaction plots normalized to observed/expected between a 1 Mb region surrounding the bait site on chromosome 2 and the rest of chromosome 2, +/-4OHT. Arrows represent frequently cut AsiSI sites. Data adjacent to the bait site along the main diagonal (grey box) has been omitted (11000000 to 16000000 Mb). Numbers (1-3) indicate three intrachromosomal sites analyzed in **Fig. 3b**. Bottom: Normalized translocation frequency (translocations per 1,000 events in the dataset) between bait and chromosome 2 loci following damage. **c**, Log-log correlation plot of translocation frequency versus Hi-C interaction frequency (+4OHT) for the 19 most common prey sites on chromosome 2. Pearson correlation (“*r*”) is shown. Points within 1 Mb of the bait site have been excluded.

Next, we sought to visualize clustering of individual DSBs within a single chromosome (Chr 2). We analyzed Hi-C interactions (normalized to observed/expected) between the area surrounding a frequently cleaved DSB on chromosome 2 and 1 Mb bins spanning the rest of chromosome 2. For differential interaction plots, blue bars above the axis indicate strengthened interactions following damage, while red bars below the axis represent a decrease in interaction frequency (**Fig. 2b**). Concordant with APA analysis (**Fig. 2a**), we found that following damage this reference site (gray) interacted with other DSBs in *cis* (arrows) with frequencies significantly higher than in the absence of damage (**Fig. 2b, top**). Next, we assessed the impact of DSB clustering in *cis* (**Fig. 2b**) on intrachromosomal translocations. Indeed, a significant fraction of oncogenic translocations takes place between loci on the same chromosome^29-31^. Thus, we performed high-throughput genome-wide translocation sequencing (HTGTS) to test whether genomic reorganization following damage influences chromosome rearrangements. HTGTS identifies translocation events between a fixed “bait” DSB and “prey” sites throughout the genome (**Extended Data Fig. 4a**)^32^. To explore how damage-induced chromatin reorganization influences aberrant rearrangements, we used the same AsiSI reference site on chromosome 2 as the bait site. HTGTS analyses revealed that the loci of heightened interactions corresponded to sites of frequent translocations with the bait (**Fig. 2b, bottom**).

We then assessed the correlation between contact frequency (+4OHT) and translocation frequency across chromosome 2. We observed that chromatin contact frequency predicted translocation occurrence with a significant Pearson correlation coefficient, *r*, of 0.6490 (**Fig. 2c**). Thus, DNA repair 23 domains are sites where translocations can occur.

### ARP2/3-mediated DSB clustering facilitates genomic rearrangements

Given that approximately 100 AsiSI sites are efficiently cut upon induction of AsiSI-ER, we predicted that most recurrent translocations would take place between active AsiSI loci. Indeed, more than 80% of prey originated from within 500 bp of an AsiSI site (**Fig. 3a**), whereas approximately 15% of translocations occurred 10 kb - 100 Mb away from an AsiSI motif. The distribution of prey sites in U2OS cells revealed similar classes (proximal, distal) of translocations (**Extended Data Fig. 4b**). Translocations between AsiSI-proximal sites and the bait are recurrent, with variable levels of end-resection, mostly under 500 bp. Recurrent translocations are not identical at the nucleotide level as PCR duplicates are filtered by the HTGTS pipeline. Translocations to distal prey are primarily unique translocations (**Extended Data Fig. 4c**) and might involve spontaneous, physiological DSBs forming at sites of intrinsic genome fragility, including R-loops, G4 quadruplex, stalled replication forks, and active transcription^33^. We showed that frequently cleaved AsiSI sites are within transcriptionally active regions (**Extended Data Fig. 2g**). Similarly, translocations originating from loci proximal to AsiSI sites were highly enriched in promoter sequences (**Extended Data Fig. 4d**) and located in transcriptionally active areas, as seen by GSEA (**Extended Data Fig. 4e**). In contrast, translocations originating from regions distal to AsiSI sites occurred throughout the genome (**Extended Data Fig. 4d**).

**Figure 3.**
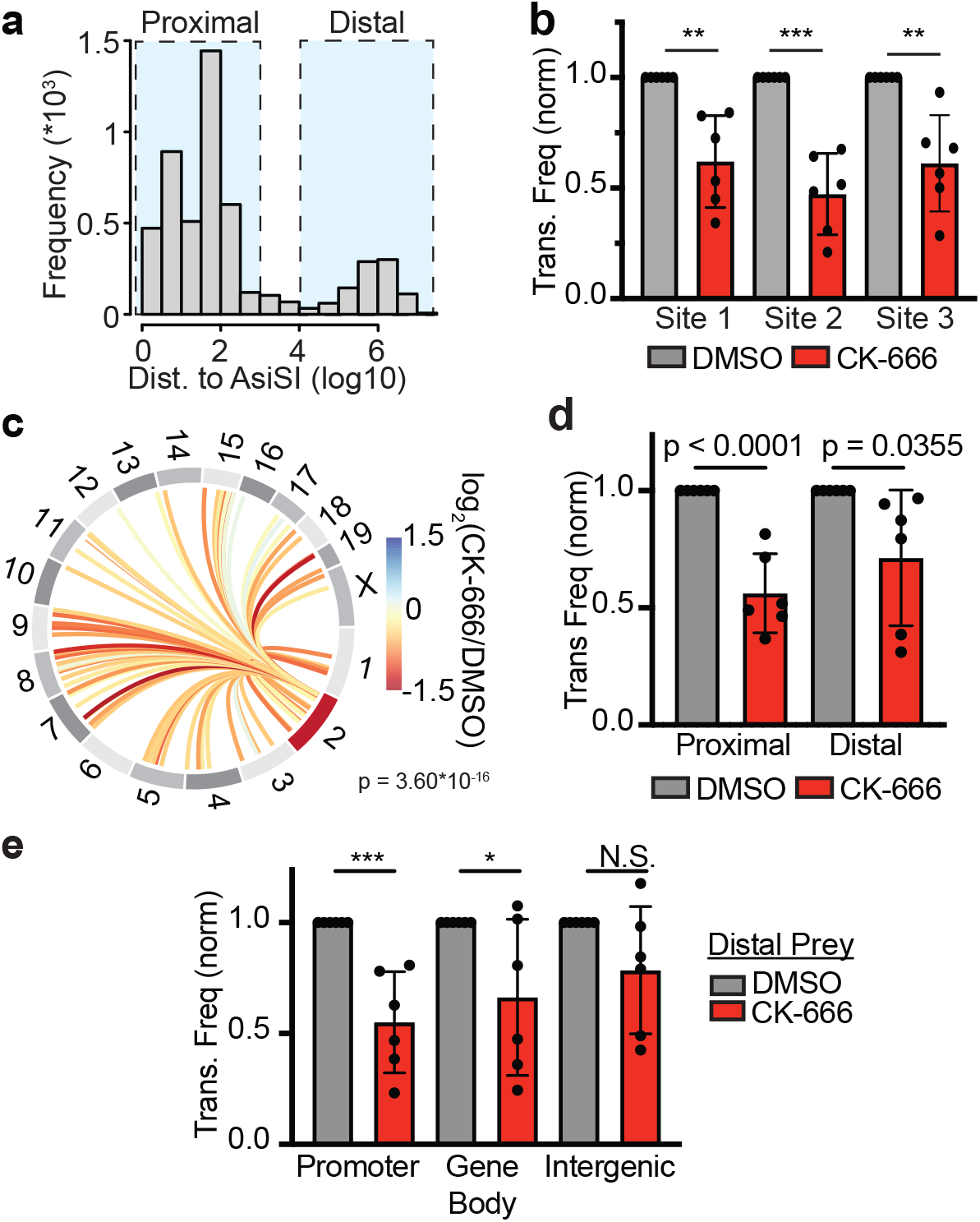
ARP2/3-mediated clustering facilitates chromosomal translocations. **a**, Plot of all translocations as a function of their distance to the nearest AsiSI motif. Data is divided into proximal (<500 bp of an AsiSI site) and distal (>10 kb from an AsiSI site) prey **b**, Normalized translocation frequency at three sites on chromosome 2 (1 = 31,900,000–32,000,000 bp; 2 = 153,600,000–153,700,000 bp; 3 = 156,100,000–156,200,000 bp) in WT MEF AsiSI cells +/-100 μM CK-666. *P* calculated by Student’s two-tailed t-test. Mean and standard deviation. **c**, Circos plot visualizing differential normalized translocation frequencies genome-wide following damage in the presence or absence of ARP2/3 inhibitor, CK-666 (100 μM) at binned loci that had ≥10 translocation events. Connecting lines are colored according to the log_2_ fold change following damage between +/-CK-666 populations. Chromosome 2 (red) contains the bait site. *P* = 3.60 * 10^−16^, Wilcoxon test. **d**, Normalized translocation frequencies to proximal (<500 bp from AsiSI site) and distal (>10 kb from AsiSI site) loci in the presence and absence of 100 μM CK-666. *P* calculated by Student’s two-tailed t-test. Mean and standard deviation. **e**, Normalized translocation frequencies for distal prey in promoter, gene body, and intergenic regions in the presence or absence of CK-666. *P* calculated by Student’s two-tailed *t*-test. Mean and standard deviation.

ARP2/3 facilitates distant DSB-DSB interactions (**Fig. 2a**) and promotes clustering of repair foci^2^. However, it is not known whether nuclear actin dynamics impact the frequency of chromosome translocations. To establish formally that increased interactions between DSBs drives chromosome rearrangements, we assessed the impact of ARP2/3 inhibition on translocations genome-wide, using HTGTS. Translocations were monitored six hours after DSB induction in control cells and in cells treated with CK-666. CK-666 significantly decreased the normalized frequency (see methods) of both intra-chromosomal (chromosome 2), p < 0.01 for the three binned loci (**Fig. 3b**) and inter-chromosomal translocations, p = 3.6.10^−16^ (**Fig. 3c**). Furthermore, the fold decrease (CK-666/DMSO) in normalized, AsiSI-proximal translocation frequency was comparable for intra- and inter-chromosomal translocations (**Extended Data Fig. 4f**). This establishes that ARP2/3-dependent actin nucleation is a driving force for chromosome rearrangements.

We then asked how DSB mobility affected translocations to spontaneous DSBs. The frequency of translocations to distal DSBs was significantly decreased following treatment with ARP2/3 inhibitor, albeit to a lesser extent than the frequency of translocations to proximal DSBs (**Fig. 3d**). Thus, a smaller fraction of physiologic translocations is driven by nuclear actin polymerization. The propensity to translocate in the presence of CK-666 could reflect intrinsic properties of the prey loci, including their transcriptional activity. Therefore, we examined the impact of ARP2/3 inhibition on translocations originating from DSBs in promoter, gene body, and intergenic regions (**Fig. 3e**). ARP2/3 inhibition significantly reduced the frequency of translocations arising from spontaneous DSBs in promoter regions, a decrease that mirrored the effect of CK-666 on recurrent, experimentally-induced (AsiSI-AsiSI) rearrangements (compare **Fig. 3d** and **Fig. 3e**). ARP2/3 inhibition also modestly decreased translocations initiating from spontaneous DSBs in gene bodies. In contrast, ARP2/3 inhibition did not have a statistically significant impact on translocations arising from intergenic loci.

DNA sequences at translocation junctions provide further insight into the repair mechanisms driving rearrangements^34,35^. Specifically, the presence of limited microhomology (MH) suggests repair by alternative end-joining (alt-EJ) whereas blunt-end ligation indicates repair by classical NHEJ (c-NHEJ)^36^. We found that only 18% of junctions resulted from blunt-end ligation (**Extended Data Fig. 5a**). In contrast, 69% of junctions harbored microhomologies (MH), emphasizing the importance of alt-EJ in mediating pathologic repair (**Extended Data Fig. 5a**). Unexpectedly, we observed that 13% of translocations contained additional short insertions. These complex rearrangements did not arise from direct ligation of blunt ends or from annealing of staggered DNA ends between the bait and prey chromosomes (**Extended Data Fig. 5a**). Of note, larger inserts (> 30 bp) were not detected due to the limits of HTGTS analysis, suggesting that insertions are more frequent than we report. To explore the origins of insertion events, we mapped inserts (20 bp – 30 bp) for all proximal reads. The majority of inserts (> 80%) mapped to the vicinity of prey loci, often on the antiparallel strand (**Extended Data Fig. 5b,c**). This suggests that an intermediate step, possibly a transient invasion or annealing event, took place prior to the ligation that gave rise to a stable translocation.

### Distinct roles for BRCA1 in regulating translocations

ARP2/3 clusters spontaneous and endonuclease-generated DSBs into HDR domains, where translocations occur. Inhibition of Mre11-dependent resection, the initial step of HDR, impairs DSB mobility and ARP2/3-mediated clustering^2,15,37^, highlighting a role for resection in the formation of repair domains. In addition to promoting end-resection at HDR breaks, BRCA1 safe-guards against chromosome translocations, as evidenced by the accumulation of genomic rearrangements in BRCA1-deficient tumors^38-40^. Therefore, we sought to examine the impact of BRCA1 loss on chromosome mobility and translocations. We used BRCA1^Δ11^ AsiSI MEF cells with a truncated BRCA1 that lacks exon 11, impairing DSB resection^16,41^. We first confirmed that cleavage at the chromosome 2 bait site was comparable in WT and BRCA1^Δ11^ cells (**Extended Data Fig. 6a**), then performed HTGTS in BRCA1-deficient MEFs. The frequency of recurrent translocations, both intra-chromosomal (chromosome 2), p < 0.001 (**Fig. 4a**) and inter-chromosomal, p = 9.58*10^−24^ (**Fig. 4b**) between AsiSI-AsiSI DSBs was markedly decreased in BRCA1^Δ11^ MEFs. This finding is consistent with the role of BRCA1 in promoting end-resection, which occurs upstream of ARP2/3 activity. Indeed, inhibition of ARP2/3 in BRCA1-deficient cells did not further reduce translocation frequency (**Fig. 4a**). We next evaluated the link between DSB resection and mobility by performing live-cell imaging of BRCA1^Δ11^ MEFs. We found that mean square displacement (MSD) of NBS1 repair foci, which is recruited prior to resection of DSBs, was substantially lower in BRCA1-deficient cells as compared to WT (**Fig. 4c**). These findings further strengthen the idea that recurrent translocations are facilitated within HDR domains, the site of BRCA1 action.

**Figure 4.**
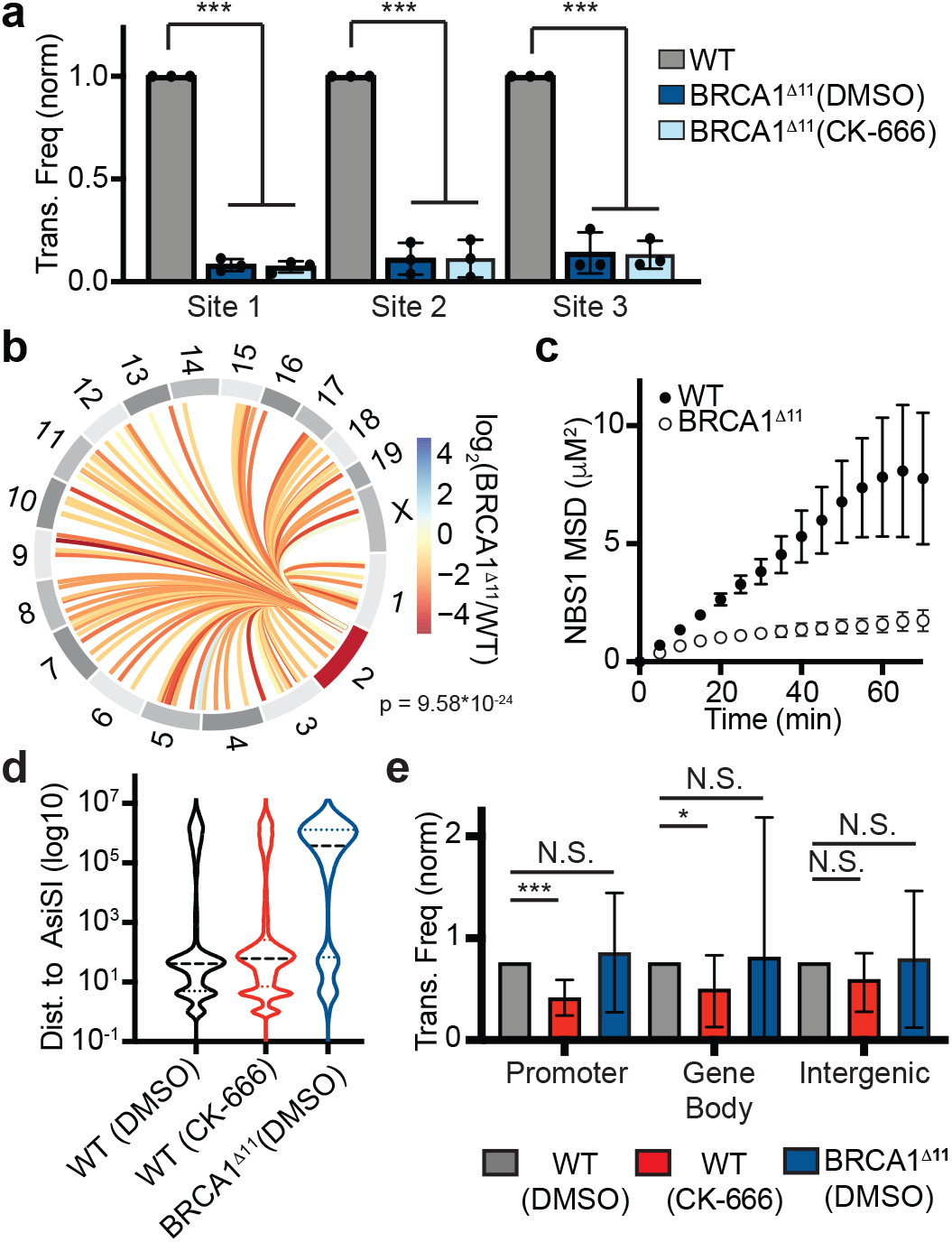
Distinct roles for BRCA1 in regulating translocations. a Figure 4| Distinct roles for BRCA1 in regulating translocations. **a**, Normalized translocation frequencies at three binned sites on chromosome 2 (1 = 31,900,000–32,000,000 bp; 2 = 153,600,000–153,700,000 bp; 3 = 156,100,000–156,200,000 bp) in WT and BRCA1^Δ11^ MEF AsiSI cells +/-100 μM CK-666. Columns are normalized to frequency of translocations in WT cells in the same biological replicate. *P* calculated by one-way ANOVA Tukey’s multiple comparisons. Mean and standard deviation. **b**, Circos plot showing differential normalized translocation frequencies following damage in BRCA1^Δ11^ cells compared to WT. Connecting lines are colored according to the log_2_ fold change between WT and BRCA1-deficient cell types following damage. Chromosome 2 (red) contains the bait site. *P* = 9.58*10^−24^, Wilcoxon test. **c**, Mean-squared displacement of NBS1-GFP foci in WT and BRCA1^Δ11^ cells treated with 0.5 μg/ml NCS. *n* > 195 foci in > 12 nuclei. **d**, Violin plot displaying the distribution of translocating prey as a function of the distance to the nearest AsiSI site in WT cells +/-CK-666 and BRCA1^Δ11^ cells (dashed line = median, dotted lines = quartiles, n > 4 biological replicates). **e**, Normalized translocation frequencies for distal prey in promoter, gene body, and intergenic categories in WT cells (+/-CK-666) and BRCA1^Δ11^ cells. *P* calculated with Student’s two-tailed t-test. Mean and standard deviation.

Whole genome sequencing of tumors harboring BRCA1 mutations has revealed rearrangement signatures thought to be the consequence of BRCA1’s role in suppressing endogenous genome instability^38,40^. Therefore, we next examined how BRCA1-deficiency might specifically affect spontaneous translocations by analyzing the distribution of prey as a function of distance to the nearest AsiSI site (**Fig. 4d**). Strikingly, translocations in BRCA1^Δ11^ cells occurred more frequently between the bait and distal DSBs than in WT cells +/-CK-666 (**Fig. 4d,e; Extended Data Fig. 6b-d**), indicating a distinct role of BRCA1 in preventing translocations. Furthermore, analysis of the cumulative frequency of prey distribution in WT and BRCA1^Δ11^ cells as a function of the distance to AsiSI sites revealed significantly different distributions (**Extended Data Fig. 6b**). This increase in translocations to distant loci could manifest from replication fork collapse or transcription-related stress as both processes are resolved by intact BRCA1^42,43^ (**Extended Data Fig. 6e**).

## Conclusions

DNA damage triggers local signaling to facilitate repair reactions at DNA lesions^44^, subsequent checkpoint activation yielding long-range histone and post-translational modifications^45^, and ARP2/3-mediated DSB mobility^2,4^. Together, these events promote the formation of HDR domains. Our studies provide insights into the coordinated, multiscale reorganization of the 3D genome leading to the formation of these domains. First, we observe local strengthening of insulation boundaries at DSBs^25,26^ (**Fig. 1c; Extended Data Fig. 7, 1**) and increased chromatin loop extrusion at CTCF boundaries^46-48^ (**Extended Data Fig. 2d-f**), both possibly the result of increased cohesin loading. Second, we provide a genomic view of DSB clustering (**Fig. 2a; Extended Data Fig. 7, 2**). Finally, we document DNA damage-dependent, genome-wide changes in compartmentalization that can be quantified as B to A compartment flips (**Fig 1a, b, Extended Data Fig. 7, 3**). Notably, damage-induced, long-range reorganization, such as clustering of DSBs and compartment flips, is facilitated in part by ARP2/3-dependent forces, whereas local changes in insulation and loop extrusion are not. These data are consistent with a model in which chromatin accessibility following damage is favored within A compartments. In turn, this facilitates DSB clustering and the generation of HDR domains, while repair activity surrounding individual DSBs is restricted by enhanced insulation.

Using high throughput translocation assays (HTGTS), we show that ARP2/3- and resection-mediated formation of HDR domains increases the risk of chromosomal translocations while facilitating homologous recombination^2^. The increased contact frequency revealed by Hi-C is not just due to rearrangements as translocation events are much more rare. We confirm that HTGTS is a powerful method for identifying translocations to naturally unstable loci, establishing that ARP2/3’s impact on chromosomal rearrangements is not limited to restriction endonuclease-generated DSBs but is also relevant for physiological damage. Nevertheless, the partial impact of ARP2/3 inhibition points to additional mechanisms for DSB clustering and pathogenic translocations, which may include different actin nucleators^15,49^, alternate cytoskeleton proteins^4,50^, and phase-separated boundaries^51,52^.

Chromosome translocations require an end-joining step^53^ (data not shown). Here we establish that clustering of resected DNA ends arising from transcriptionally active loci is also critical for translocations. We thus propose that translocations are generated by a two-step process. First, actin nucleators (ARP2/3) and resection machinery (BRCA1) bring recurrent and spontaneous DSBs harboring resected ends into close proximity (**Extended Data Fig. 7, 2**). Second, resected DNA ends are processed to be compatible with end-joining reactions or alternatively, transiently invade the prey locus capturing additional sequences prior to end-joining. This is consistent with frequent insertion events observed previously at translocation junctions^54,55^ as well as in this study (**Extended Data Fig. 5b**). Finally, we establish that while BRCA1-dependent resection facilitates DSB mobility, increasing translocations between recurrent DSBs, the tumor suppressor maintains genome integrity during DNA transactions, preventing spontaneous translocations to fragile genomic regions (**Extended Data Fig. 7**). Overall, our work highlights the delicate balance between faithful repair and misrepair at play within HDR domains and the critical roles of actin nucleators and repair proteins in achieving this balance.

## Methods

### Cell culture and drug treatment

Mouse embryonic fibroblast (MEF) and U2OS cell lines were cultured in high-glucose Dulbecco’s modified Eagle’s medium supplemented with L-glutamine, 10% fetal bovine serum, and 1% penicillin-streptomycin. ER-AsiSI MEF cell lines, including WT and BRCA1^Δ11^ cells, were developed as previously described^16^. Cells were treated with doxycycline (Sigma-Aldrich: D3072, 3 μg/mL) for 24 hours to induce AsiSI expression. 4-OHT (Sigma-Aldrich: H7904, 1 μg/mL) was added for the last 6 hours of doxycycline treatment to induce AsiSI translocation. Cells were co-treated with DMSO or 100 μM CK-666 (Sigma Aldrich: SML-006, 100 μM) and incubated at 37°C for 6 hours.

ER-AsisI U2OS cells were obtained from Dr. Gaelle Legube^13^. For cell synchronization, cells were treated with 2 mM thymidine for two 18-h intervals separated by an 11-h release in fresh medium. Following double-thymidine block, cells were released into fresh medium for 7 h (G2) or 15 h (G1). Exponentially growing or synchronized cells were treated with 300 nM 4-OHT (Sigma Aldrich, H7904) to induce damage with DMSO or 100 μM CK-666. Cells were incubated at 37°C for 24 hours.

### Hi-C

Chromosome conformation capture experiments were performed as previously described^33^ with some modifications. Briefly, 5 million cells/library were crosslinked with 1% formaldehyde and lysed. After digesting chromatin with 400 units of DpnII overnight, DNA ends were labeled with biotinylated dATP using 50 units Klenow DNA polymerase. Blunt-end ligation was performed with 50 units T4 Ligase at 16°C for 4 hours, followed by reverse crosslinking with 400 μg/ml proteinase k at 65°C overnight. DNA was purified using phenol/chloroform extraction and ethanol precipitation, and concentrated on a 30 kDa Amicon Ultra column. Biotin was removed from unligated ends in 50μl reactions using 50 units T4 DNA polymerase/5 mg DNA. Following DNA sonication (Covaris S220) and Ampure XP size fractionation to generate DNA fragments of 100-300 bp, DNA ends were repaired using 7.5U T4 DNA polymerase, 25U T4 polynucleotide kinase, and 2.5 U Klenow DNA polymerase. Libraries were enriched for ligation products by biotin pulldown with MyOne streptavidin C1 beads. To prepare for sequencing, A-tailing was performed using 15 units of Klenow DNA polymerase (3’-5’ exo-) and Illumina TruSeq DNA LT kit Set A indexed adapters were ligated. Libraries were amplified in PCR reactions for 5-7 cycles and subjected to Ampure XP size selection prior to sequencing on an Illumina HiSeq 4000 machine using the Paired End 50 bp module. Two biological replicates were performed for each condition.

### Hi-C Analysis

Paired-end 50bp reads were processed using the distiller pipeline^56^. First, reads from MEF and U2OS libraries were mapped to mm10 and hg19 reference genomes, respectively, using BWA-MEM in single sided mode (-SP). Alignments were then parsed, classified, and filtered using pairtools^56^. The resulting valid pairs included uniquely mapped and rescued pairs with a minimum mapping quality of 30. Valid pairs were aggregated into binned contact matrices and kept as multi-resolution cooler files^57^ for subsequent analyses. Where indicated, paired reads from replicate libraries were pooled prior to filtering for PCR duplicates. All Hi-C contact matrices were normalized by iterative correction^19^, excluding the first 2 diagonals. Downstream analyses were performed using cooltools version 0.3.2^58^ unless otherwise indicated, python 3.7.10, and matplotlib (Hunter, 2007). Hi-C interaction heatmaps were generated from balanced 250 kb resolution coolerfiles using cooler “show”. For heat maps, pooled and individual replicate library sets were downsampled to equal read depth.

The average contact probability (*P(s)*) as a function of genomic distance was calculated using “compute-expected” from cooltools version 0.4.0^59^. The “diagsum” function was applied to balanced data binned at 1 kb to compute expected, which was then parsed into log-spaced bins of genomic distance using “logbin-expected”. The rate of contact frequency decay as genomic distance increases, the *P(s)* derivative, was determined using “combine_binned_expected” and provides a highly informative representation of Hi-C data.

Active and inactive chromatin compartments were assessed based on eigenvector decomposition of observed/expected *cis* contact matrices binned at 250 kb resolution using the cooltools “call-compartments” function. In this case, the first eigenvector, EV1, positively correlated with gene density and assignment of A or B compartment identity was based on high or low gene density, respectively. Saddle plots for *cis* interactions were generated using cooltools “compute-saddle”. For each library, ranked EV1 values were binned into 30 quantiles and observed/expected interactions were plotted. Saddle strength was quantified by comparing the average interaction frequency of each AA or BB quantile bin or bins to the analogous AB and BA bins (effectually (AB+BA)/2).

Average observed/expected Hi-C interaction frequencies at subsets of genomic loci were determined using the cooltools “snipping” function. To examine DSB clustering, all pairwise *cis* interactions between bin-aligned Top97-digested AsiS1 sites were aggregated at 25 kb resolution with a 2 Mb flanking window. The DSB cluster enrichment score was calculated by taking the ratio of the average Hi-C interaction frequency in the 5×5 central bins (25 kb radius) and the average interaction frequency of the remaining bins (125 kb – 1 Mb radius). Loop extrusion was explored by examining aggregate Hi-C interactions at CTCF sites. CTCF positions were determined using a previously published CTCF ChIP-seq dataset from MEFs^60^ (sample GSM2635593). Peaks were called using MACS3 (https://github.com/macs3-project/MACS) with the default “callpeak” parameters and candidate CTCF motifs, generated in HOMER^61^ using a published vertebrate consensus^62^ within 200 bp of these peaks were selected. For pileups, top CTCF sites (13927 total) were flipped based on the direction of the consensus motif and aggregated at 5 kb bin resolution with a 100 kb flanking window. Loop aggregate plots were generated by considering all possible pairwise combinations of convergent CTCF sites on cis chromosomes with a genomic distance of 20-1000 kb (64044 possible loops). Loop scores were calculated by taking the ratio of the average Hi-C interaction frequency in the 5×5 central bins (25 kb radius) and the average interaction frequency of the remaining bins (25-100 kb radius).

### High-throughput Genome-wide Translocation Sequencing

HTGTS was performed as previously described^32^. Briefly, genomic DNA was collected using phenol/chloroform extraction, sonicated (Covaris S220), and amplified using biotin (MEF: 5’ Bio-TGGAGAGCGATGAACTGGATC 3’; U2OS: 5’-Bio-GCCGACCAATAGCATGGCG-3’) and nested (MEF: 5’-NNNNNN***Barcode**CGAAAACAGGATCCCGCAGC*-3’; U2OS: 5’-***Barcode***ACTGCGGCTGCATCCAATC-3’) primers targeting chromosome 2 (MEFs, chr2: 13271321) and chromosome 9 (U2OS, chr9: 130693175). For the nested primer in MEF experiments, random nucleotides were added before the barcode to increase library diversity. Sequencing was performed on an Illumina MiSeq sequencer.

### High-throughput Genome-wide Translocation Analysis

Burrows-Wheeler Aligner was used to align sequences to the mm10 (MEF) or hg19 (human) genomes. Using established pipelines (https://github.com/robinmeyers/transloc_pipeline), reads were filtered with the default parameters. All reads had good mapping quality (mapping quality >30). For translocation frequency, final reads were binned by 100 kb windows genome-wide. For each experiment, the number of reads in each window was normalized to the corresponding number of bait-only sequences obtained from the pipeline allowing us to compare translocation frequency between libraries. Genome coordinates of prey sequences were annotated using R package ChIPseeker^63^, which retrieved the location of each prey sequence (Promoter, Gene Body or Intergenic Region). Microhomology (MH) analysis was performed as previously described^35^. MH was defined as the overlapping homologous sequence between the bait and the prey site.

### Live-cell Imaging

MEF cells were transfected with plasmids expressing NBS1-GFP using Neon Transfection System (1350 V, 30 ms, 1 pulse). Cells were cultured on 35-mm glass bottom microwell dishes (MatTek) and damaged with 0.5 μg/ml NCS (Sigma N9162) for 60 minutes at 37 °C. Following two washes with PBS, cells were allowed to recover for 10 hours before imaging. Imaging was performed on an A1RMP confocal microscope (Nikon Instruments), on a TiE Eclipse stand equipped with a 60×/1.49 Apo-TIRF oil-immersion objective lens, an automated XY stage, stage-mounted piezoelectric focus drive, and a heated, humidified stage top chamber with 5% CO2 atmosphere. Z series were collected at 0.4-μm intervals throughout the entire nucleus every 5 min for 1 hour. Focus was maintained by the Perfect Focus System (Nikon). Mean-squared displacement analysis was performed as previously described^2^.

### Immunohistochemistry and quantification of γH2AX foci

Fixed-cell imaging experiments were performed as previously described^2^. Briefly, U2OS cells were cultured on 8-well chamber slides and treated with 0.5 μg/ml NCS for 60 minutes at 37°C. Following two washes, cells were incubated at 37 °C to allow formation of **γ**H2AX foci in the presence of DMSO or 100 μM CK-666. Cells were fixed with 4% PFA (pH 7.4) and permeabilized with 0.1% PBS-Trition X-100. Cells then were treated with primary antibody (γH2AX, EMD Millipore: 05-636, 1/500) at 4 °C overnight and secondary antibody (Alexa 488 conjugated goat anti-mouse Ig (Abcam: ab150113, 1/1,000)) for 1 hour at room temperature. Cells were imaged under 40x magnification using a Zeiss Axio Imager Z2 microscope, equipped with a CoolCube1 camera (Carl Zeiss). Foci counting was performed using automated MetaCyte software (Metasystems, version 3.10.6).

### Quantification of AsiSI-induced DSBs

END-seq experiments and spike-in assays were performed as previously described^16,64^. AsiSI cutting efficiency at specific sites was measured by quantitative polymerase chain reaction (qPCR)^17^ using delta-delta Ct to compare samples +/-4OHT. Primers used in U2OS cells are as follows. Site 1: 5’-GTCCCTCGAAGGGAGCAC-3’, 5’-CCGACTTTGCTGTGTGACC-3’; Site 2: 5’-CCGCCAGAAAGTTTCCTAGA-3’, 5’-CTCACCCTTGCAGCACTTG-3’. Primers used in MEF cells are as follows. Bait: 5’-TGGAGAGCGATGAACTGGATC-3’, 5’-TGGCCGGATTTTGTGTGC-3’. Ct was normalized for DNA content using primers distant from any AsiSI motifs (No DSB). In U2OS cells: 5’-ATTGGGTATCTGCGTCTAGTGAGG-3’, 5’-GACTCAATTACATCCCTGCAGCT-3’. In MEF cells: 5’-GGACAATGACCGCGTGTTTT-3’, 5’-AACAGCAGGCGCTCTATACC-3’.

### Gene Set Enrichment Analysis

GSEAPreranked was used to assess the enrichment of the frequently cut AsiSI sites in transcriptionally active regions. The transcriptional profile of MEF cell line was downloaded from GEO https://www.ncbi.nlm.nih.gov/geo/query/acc.cgi?acc=GSE29278^65^ to create the preranked gene list with the level of gene expression as the input. The closest gene to each AsISI site was collected to make the gene set *.gmt file. The same approach was used to assess enrichment of prey sites (HTGTS) in transcriptionally active regions.

## Data Availability

High throughput sequencing data have been deposited to Gene Expression Omnibus under accession number GSEXXXXXX.

## Acknowledgements

We thank the Molecular Cytogenetics and Molecular Pathology Shared Resources of the Herbert Irving Comprehensive Cancer Center (HICCC) as well as T. Swayne and E.L. Munteanu from the Confocal and Specialized Microscopy Shared Resource of the HICCC at Columbia University. We also thank G. Legube for the ER-AsiSI U2OS cell lines; R. Baer for BRCA1^Δ11^ MEFs used for live-cell imaging; and J. Min, G. Sidhu, and Y. Rose for comments on the manuscript. This work was supported by the following 31 NIH/NCI/NHGRI grants: F30-CA250166 (J.Z.), CA197606 (J.G.), CA174653 (J.G., R.R., and Ju.Z), and HG003143 (J.D.). The A.N. laboratory is supported by the Intramural Research Program of the NIH. J.D. is an investigator of the Howard Hughes Medical Institute.

## Author Contributions

J.G. and J.Z. conceived of the study and wrote the manuscript. J.Z. conducted the majority of experiments. J.Z. A.S., and Ju.Z. performed data analysis. J.D., R.R., S.Z., and M.G. aided with data interpretation. A.S. and J.D. helped with implementation of Hi-C protocols. B.S. aided with initial experiments. E.C. and A.N performed END-seq experiments.

## Extended Data Figure Legends

**Extended Data Figure 1.**
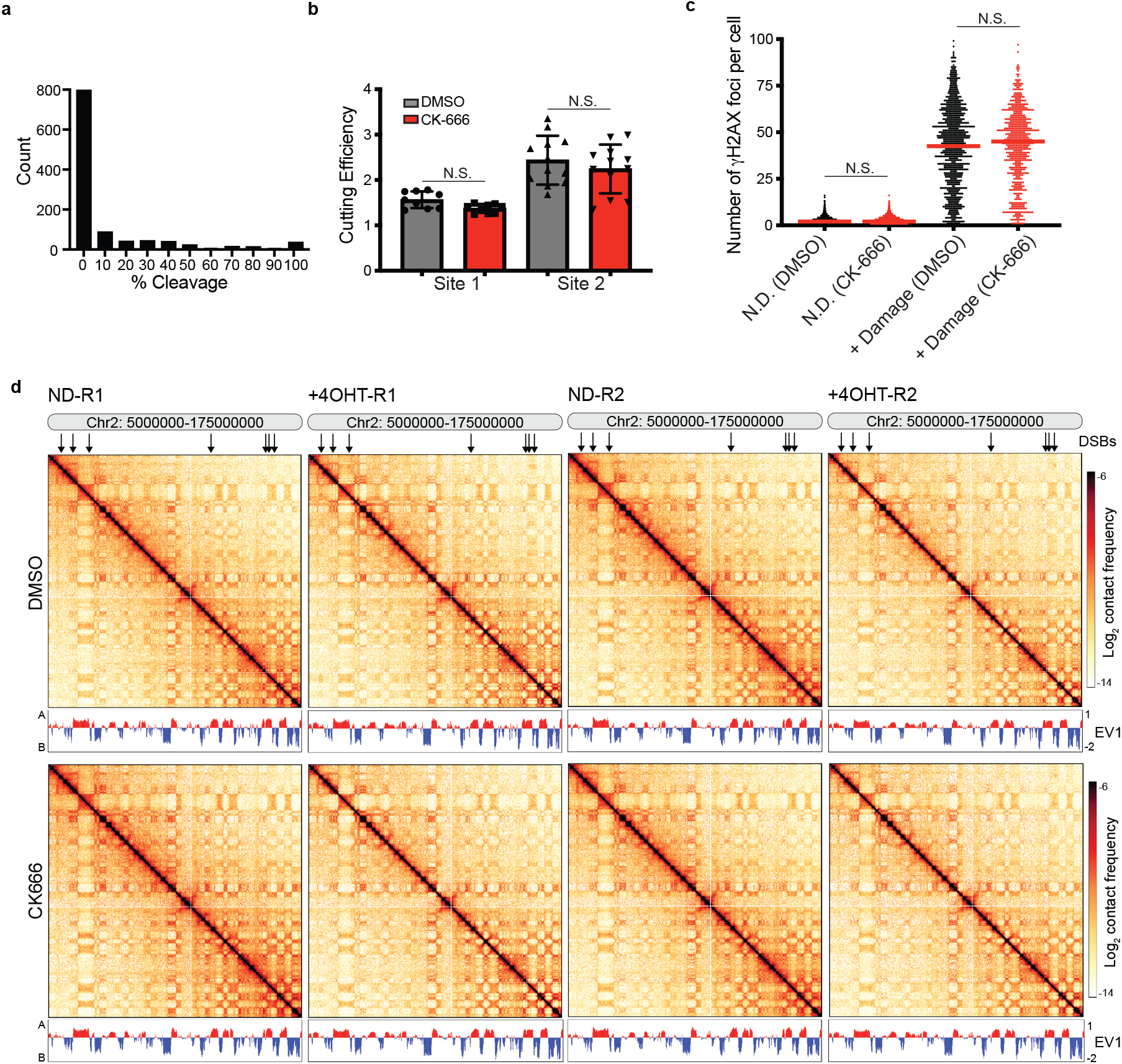
Characterization of MEF and U2OS cell lines. **a**, AsiSI restriction endonuclease cutting efficiency at all AsiSI motifs in MEFs as measured by END-seq spike-in assays. **b**, Cutting efficiency for two AsiSI sites (chr9: 130693175 and chr2: 38864106) in U2OS cells +/-100 μM CK-666. DNA was extracted from cells 4 hours following damage and % DSBs was measured using quantitative PCR amplification with primers close to the AsiSI sites, normalized to a control (uncleaved) site. Mean and standard deviation. n = 3 biological replicates with each 3 technical replicates. *P* calculated by Student’s two-tailed t-test. **c**, Quantification of γH2AX foci/cell in undamaged U2OS cells, and cells treated with 0.5 μg/ml NCS +/-CK-666. Cells were allowed to recover for two hours following damage. *P* calculated by two-sided Mann-Whitney test. Red line indicates mean. **d**, Hi-C interaction frequency maps for a region of chromosome 2 binned at 250 kb and accompanying first eigenvector traks (EV1) for *cis* interactions phased by gene density (Active/A compartment > 0). Top 97 frequently digested AsiSI sites in MEFs are indicated by arrows.

**Extended Data Figure 2.**
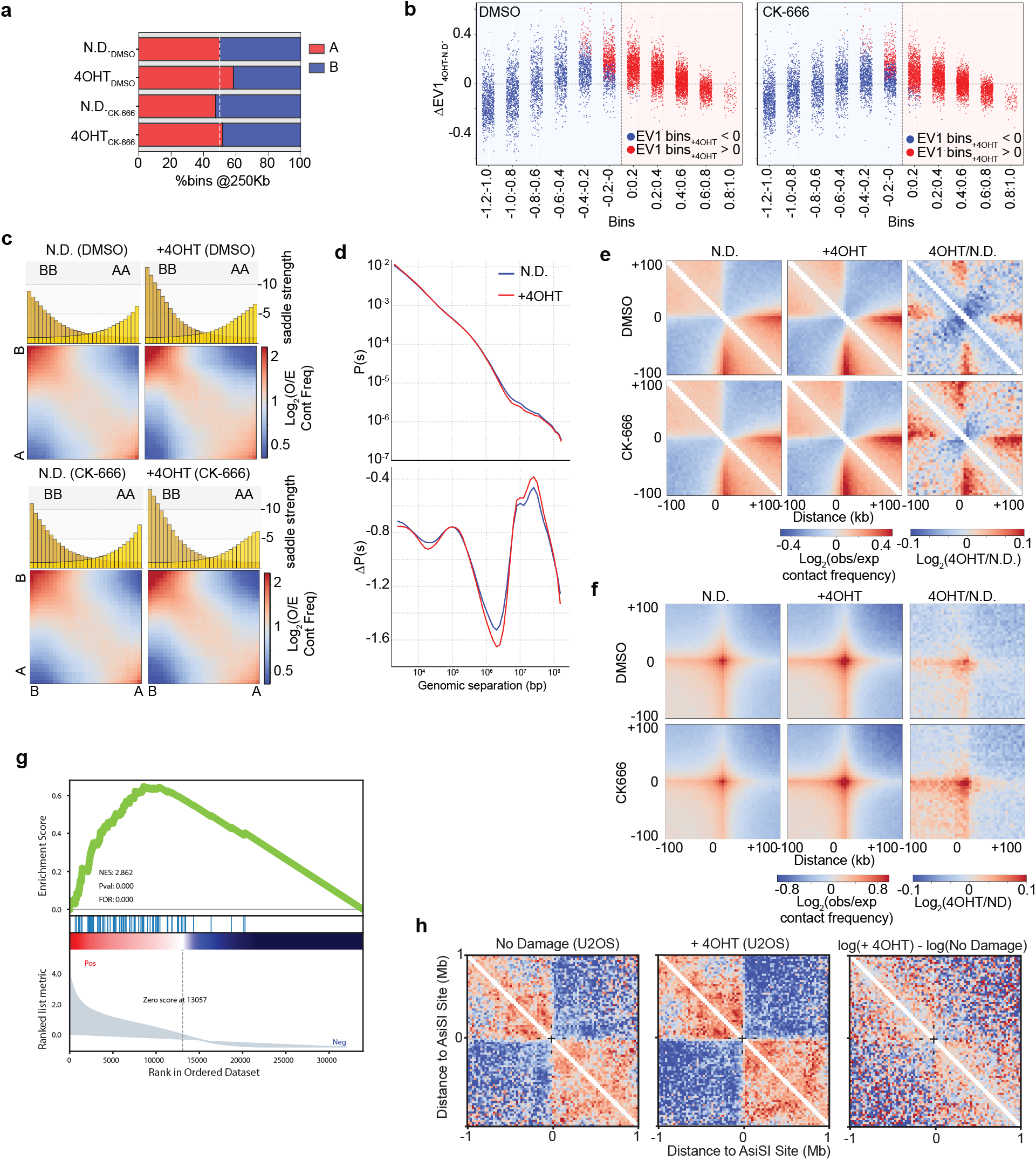
Genome reorganization following damage. **a**, Fraction of the genome (250 kb bins) classified as A (EV1>0) or B (EV1<0) compartment before and after damage +/-CK-666 (100 μM). **b**, Genome wide changes in EV1 (+4OHT vs N.D. control) plotted as a function of binned eigenvalues in the undamaged control. **c**, Saddle plots representing chromatin compartmentalization, i.e. the strength of A-A (bottom right quadrant) and B-B (top left quadrant) compartment interactions versus interactions between compartments (250 kb-binned data). Data is normalized by the expected interaction frequency based on genomic distance. Histograms along the X-axis show the distribution of saddle strength, as measured by (AA+BB)/(AB+BA). **d**, Top: Contact probability *P* plotted as a function of genomic distance *s* (*P*(*s*)) for chromosome 2 in the presence or absence of DNA damage (4OHT). Bottom: Derivative plots emphasize changes in distant gene-gene interactions. **e**, Average log_2_(observed/expected) Hi-C interaction frequency maps in the 200 kb regions flanking top CTCF sites (4052) binned at 5 kb resolution. **f**, Log_2_(observed/expected) Hi-C maps centered on all possible pairwise combinations of the top CTCF sites (11532 interactions) binned at 5 kb and averaged for a 200 kb flanking window. Log ratio interaction maps for 4OHT vs control treatments are shown. **g**, Gene set enrichment analysis (GSEA) plot (score curves) assessing the enrichment of the frequently cut AsiSI sites in transcriptionally active regions **h**, Pile-up heat maps 1 Mb surrounding the most frequently cut AsiSI sites in U2OS cells. For these experiments, no damage samples were exponentially growing, and damaged cells were synchronized in G2. Cells synchronized in G1 looked similar (data not shown).

**Extended Data Figure 3.**
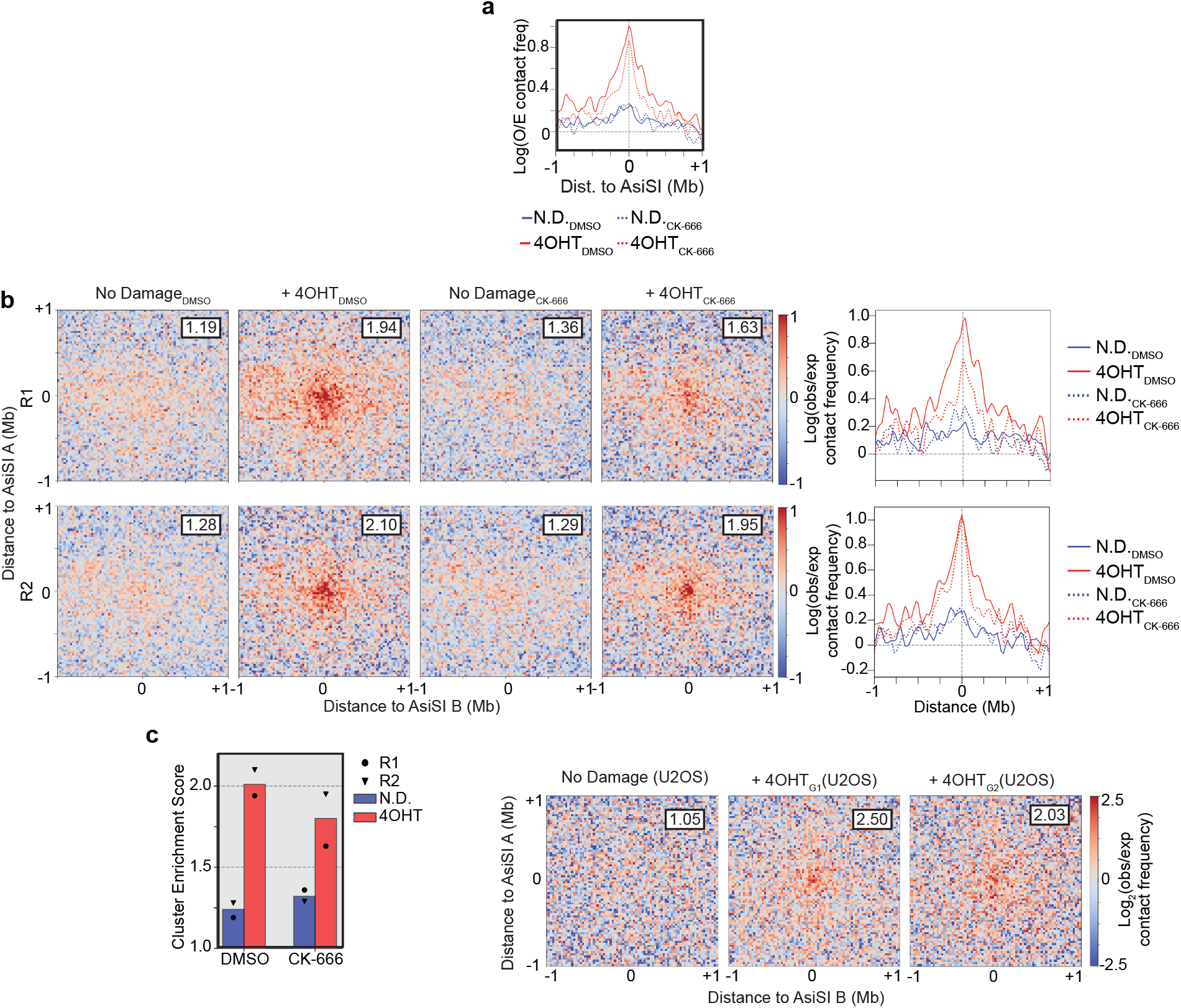
DSB clustering in mouse and human cell lines. **a**, Observed-expected contact frequencies derived from aggregate peak analysis (APA) in the presence and absence of damage +/-100 μM CK-666 (MEF cells). **b**, Left: Average Hi-C interactions centered on all possible combinations of the Top97 AsiSI digested sites *in cis* (304 interactions between damaged bins) for individual biological replicates. Data is binned at 25 kb and averaged for a 2 Mb flanking window. Log_2_(observed/expected) Hi-C maps are shown in the presence or absence of damage (4OHT) +/-CK-666. Top right corner in each aggregate peak analysis (APA) plot displays cluster enrichment score which is calculated using the ratios of the average interaction frequency of the 9 central bins (125 kb) / average interaction frequency of the outside bins (125 kb – 1 Mb). Right: Observed-expected contact frequency derived from aggregate peak analysis (APA) in the presence and absence of damage +/-100 μM CK-666 (MEF cells) for individual biological replicates. **c**, Quantification of cluster enrichment score for two biological replicates and pooled data. **d**, APA plots (2 Mb flanking window, 25 kb resolution) of interactions between the most frequently cut U2OS AsiSI sites in *cis* (155 pairs). No damage cells were exponential growing, and damaged (+4OHT) cells were synchronized in G2. Cells synchronized in G1 looked similar (data not shown).

**Extended Data Figure 4.**
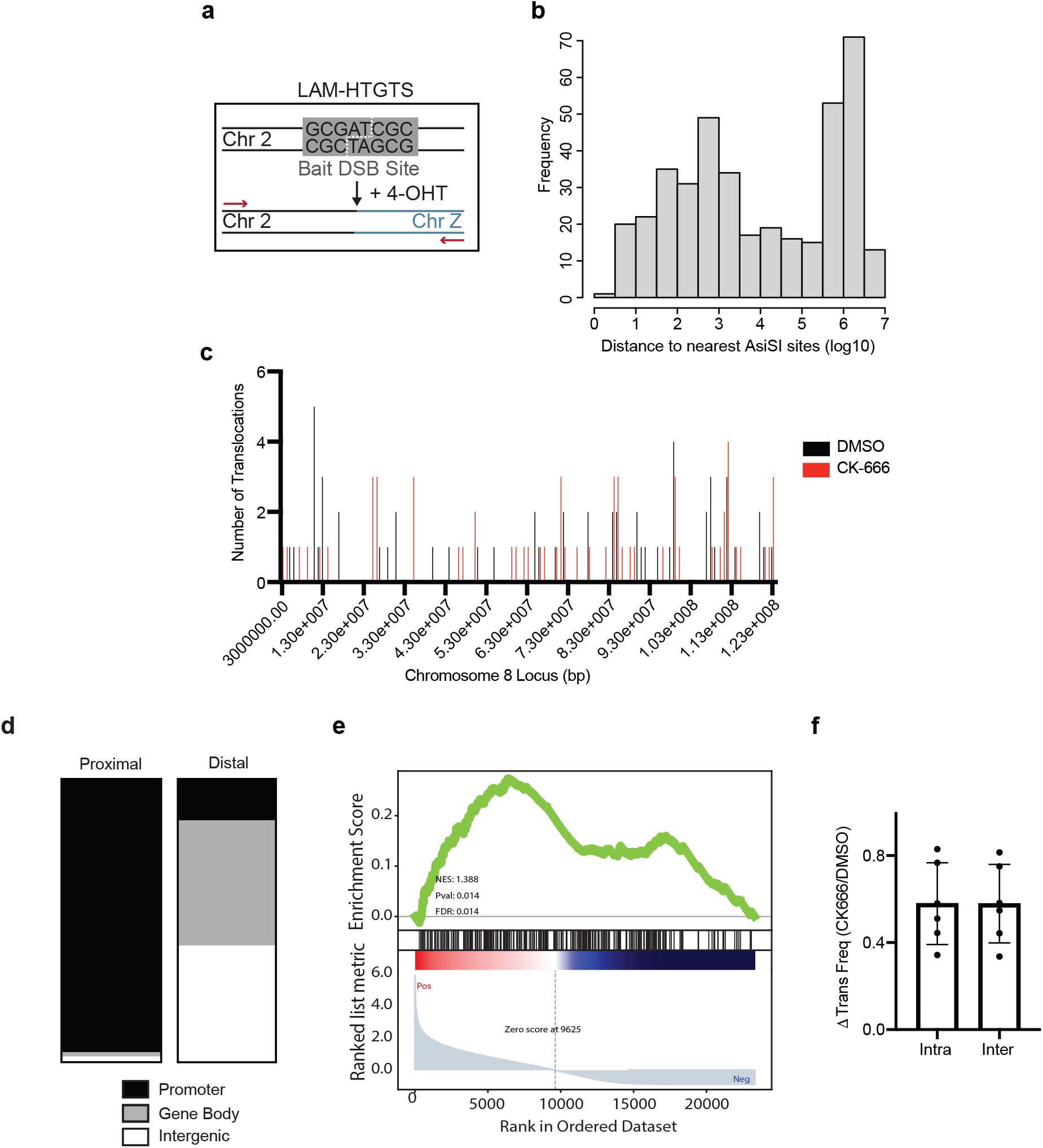
Recurrent (proximal) versus spontaneous (distal) translocations. **a**, Schematic of HTGTS experiment. The bait site is located on chromosome 2, 13271321 bp. b, Distance of translocating loci (prey) to the nearest AsiSI motif in U2OS cells. **b**, Distribution of distal prey along chromosome 8 in two individual libraries (+/-CK-666). **c**, Distribution of proximal and distal prey into promoter, gene body, and intergenic categories. **d**, Gene set enrichment analysis (GSEA) plot (score curves) assessing enrichment of translocating prey in transcriptionally active areas. **e**, Fold change in normalized translocation frequency for intra- and inter-chromosomal events. Mean and standard deviation.

**Extended Data Figure 5.**
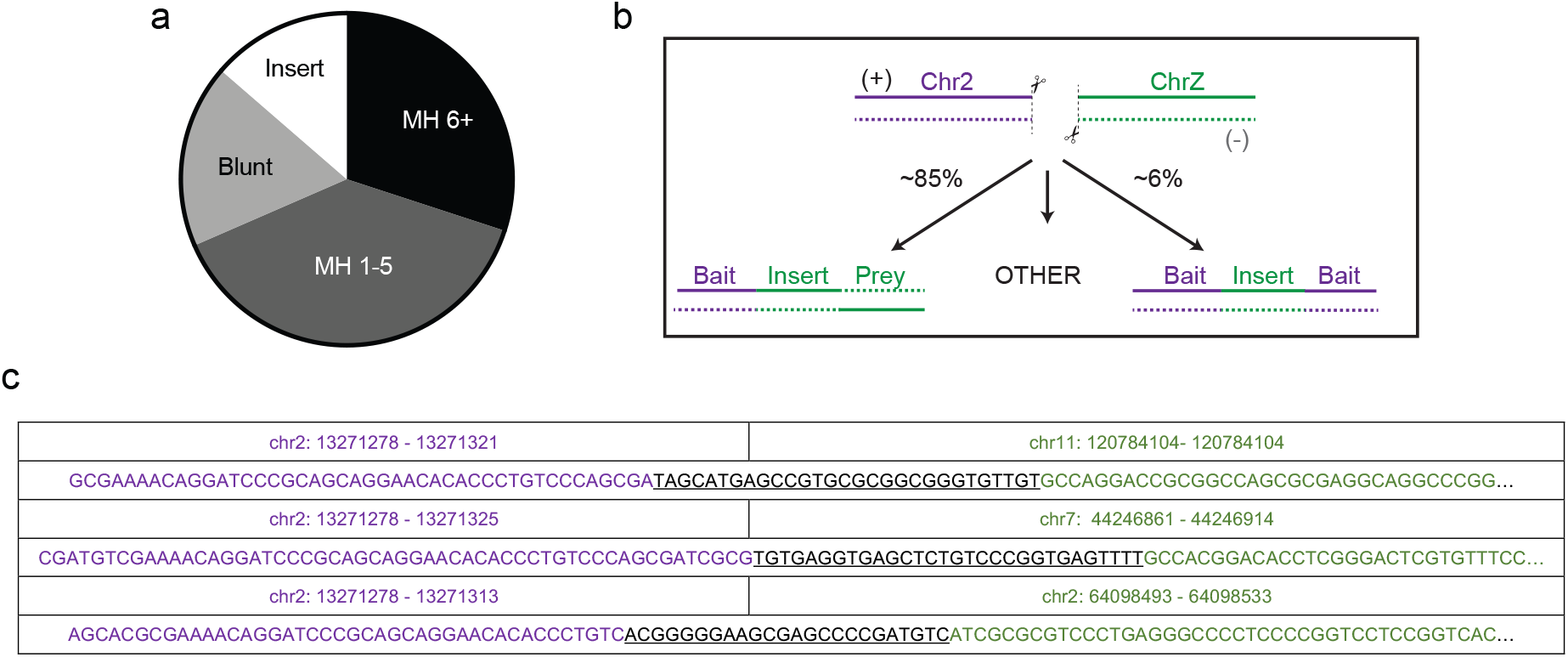
Junctional analysis of translocation events. **a**, Distribution of HTGTS prey by junctional type (MH, microhomology; blunt end ligation; insertion). **b**, Schematic representation of insertion events. **c**, Example reads that contain an insertion event (underline) in between bait (purple) and prey (green).

**Extended Data Figure 6.**
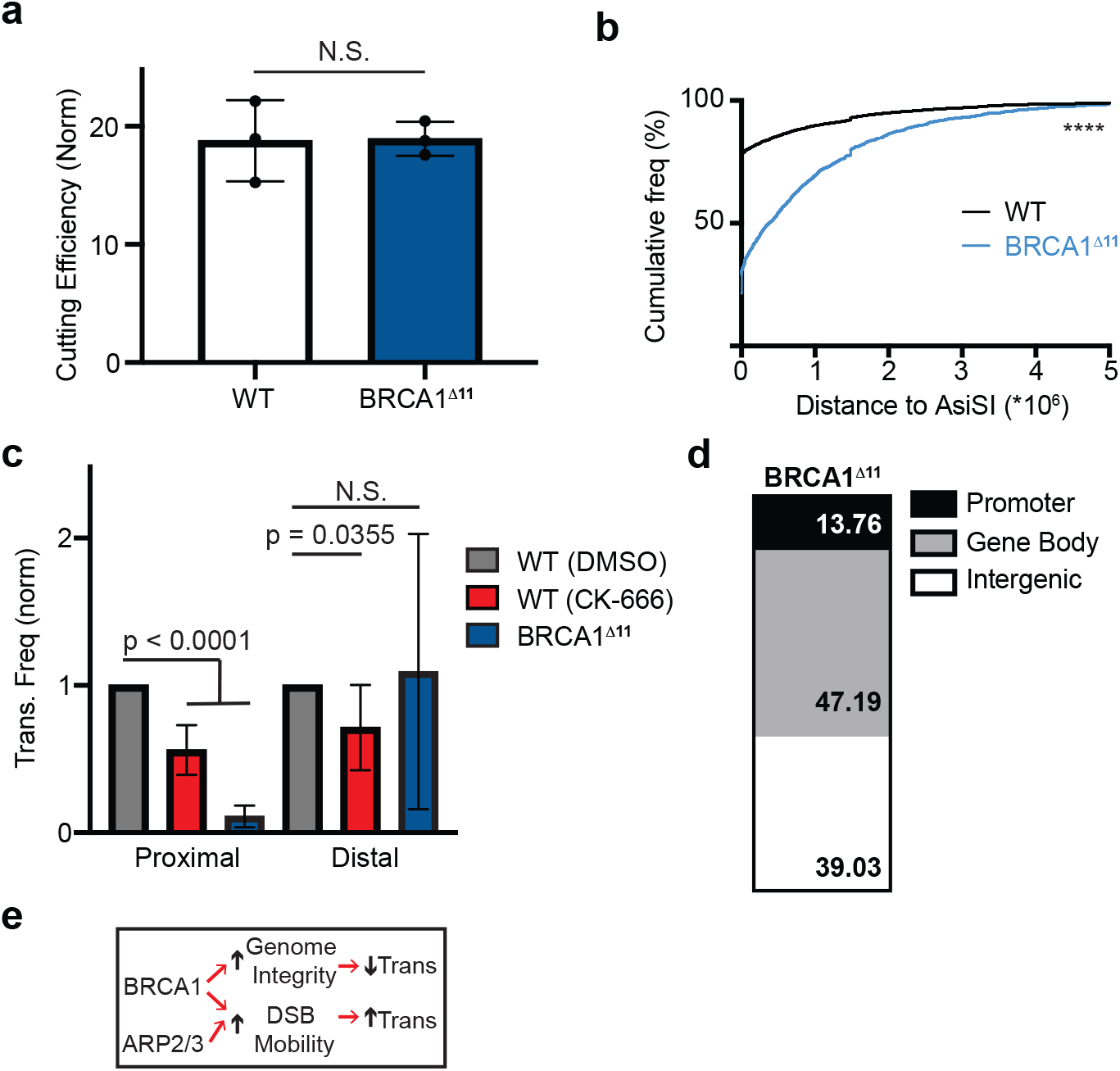
BRCA1 facilitates translocations to recurrent DSBs. **a**, Cleavage efficiency of the bait site in WT and BRCA1^Δ11^ MEF AsiSI cells. DNA was extracted from cells 6 hours following damage and % DSBs was measured using quantitative PCR amplification with primers close to the bait and normalized to a control (uncleaved) site. Mean and standard deviation. n = 3 technical replicates. Biological replicates showed comparable results. *P* calculated by Student’s two-tailed t-test. **b**, Cumulative translocation frequency as a function of distance to the nearest AsiSI site in WT and BRCA1^Δ11^ MEF AsiSI cells. *P* calculated by Kolmogorov–Smirnov test. **c**, Normalized translocation frequency to proximal and distal prey in WT and BRCA1^Δ11^ MEF AsiSI cells +/-100 μM CK-666. Columns are normalized to frequency of translocations in WT cells in the same biological replicate. *P* calculated by one-way ANOVA Tukey’s multiple comparisons. Mean and standard deviation. **d**, Distribution of prey into promoter, gene body, and intergenic categories for distal DSBs in BRCA1-deficient cells. **e**, Graphical representation of the distinct roles of BRCA1 in modulating translocations.

**Extended Data Figure 7.**
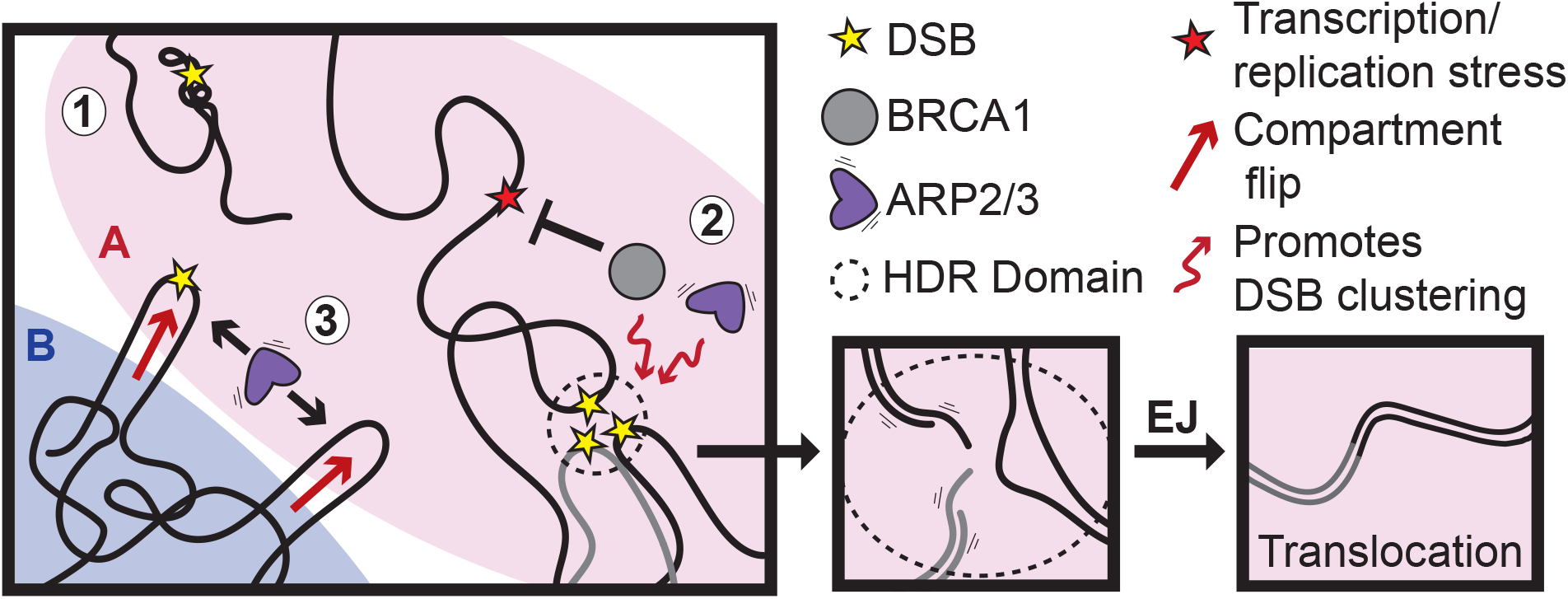
Genome reorganization following DNA damage facilitates translocations. Schematic representation of the multiscale changes in the 3D genome following damage.

